# The role of UCP3 in brown adipose tissue mitochondrial bioenergetics is complementary to that of UCP1

**DOI:** 10.1101/2020.03.24.003442

**Authors:** Christopher L. Riley, Edward M. Mills

## Abstract

Mitochondrial uncoupling protein 3 (UCP3) plays a complementary role in uncoupling protein 1 (UCP1)-dependent uncoupling in mammalian thermogenesis. Using mouse strains that lack UCP1 or UCP3, plus a UCP1/UCP3 double knockout, we have previously demonstrated that UCP3 is necessary for sustained heat production. However, how and where UCP3 is essential for heat production remains unknown. Here we use brown adipocyte mitochondria from mice lacking UCP1, UCP3, or UCP1 plus UCP3 (double knockout) we found that in the absence of UCP1, UCP3 does not act as a free fatty acid-inducible uncoupling protein; however, UCP3 is necessary for maximal GDP-sensitive respiration. We additionally confirm that the loss of UCP1 is a dominant regulator of mitochondrial respiratory chain activity, assembly, and composition in brown adipose tissue. Our data suggest without UCP1, brown adipose tissue is not a significant source of heat production, and that UCP3-dependent heat generation serves a regulatory function outside of adipose tissues. Together these findings suggest that in brown adipose tissue, UCP3’s thermogenic function is independent of its role as a *bona fide* uncoupling protein.

## Introduction

Brown adipose tissue (BAT) mediates approximately 60% of adaptive non-shivering thermogenesis in small mammals, and is an essential determinant of the arousal from torpor in hibernators[1]. BAT’s recently identified presence in a significant percentage of adult humans, along with identification of brown-like ‘beige’ adipocytes in white adipose tissue, suggest that modulation of BAT function holds great promise for the treatment of human metabolic disease associated with overnutrition [2,3].

BAT mitochondria are unique in that they oxidize substrates to generate heat, instead of ATP: this is a metabolic property that antagonizes obesity. Mitochondrial uncoupling protein 1(UCP1) is the well-established thermogenic mediator of BAT, and comprises approximately 20% of the BAT mitochondrial proteome. Stimulated by fatty acids, UCP1 ‘uncouples’ substrate oxidation from ATP synthesis by physically transporting protons from the intermembrane space back into the mitochondrial matrix[4].

While UCP1 remains unquestioned as the dominant mitochondrial uncoupling protein, the close UCP1 homolog UCP3 (57% percent sequence identity to UCP1) is also expressed in BAT, and exhibits uncoupling properties *in vitro* as well as *in vivo*[5–7]. However, little is known about the function of UCP3 in BAT mitochondrial bioenergetics. UCP3 is expressed in skin[8], white and brown adipose tissue, skeletal muscle, and heart [9]. Most physiological and biochemical studies have examined the functions of UCP3 in skeletal muscle mitochondria, where its role as a *bona fide* mitochondrial uncoupler is debated [10–12]. Its expression in skeletal muscle is highly correlated with fatty acid oxidation as well as with levels of reactive oxygen species. Nevertheless, UCP3 expression in BAT is differentially regulated, suggesting that UPC3 may function differently in BAT and skeletal muscle[13]. And in contrast to the earlier reports, Hilse *et al.* have demonstrated that the level of expression of UCP3 is two-fold higher in BAT mitochondria than in skeletal muscle.

Despite considerable interest about its role in metabolism in general, exactly what function UCP3 serves in murine BAT mitochondrial oxidation remains a mystery. We have established that UCP3 contributes significantly to the thermogenic responses to sympathomimetics and lipopolysaccharide *in vivo* [14,15]. Here we have sought to examine the bioenergetic role of UCP3 in mitochondrial function in BAT.

A recent report showed that UCP3 is expressed in BAT at higher levels that in SKM[9]. In combination with our findings *in vivo*, we hypothesized that UCP3 acts as a traditional uncoupler in BAT. The generation of a UCP1/UCP3 double knockout allowed us to characterize BAT mitochondrial uncoupling in the absence of UCP1 in BAT. Using this double knockout model, along with single UCP1 and UCP3 null mice, we found that the loss of UCP3 in brown adipose tissue decreases GDP-sensitive mitochondrial respiration in a UCP1-dependent manner, but does not significantly regulate fatty acid-inducible uncoupling. We also confirmed previous reports that UCP1 is necessary for maximal and inducible respiration, and controls mitochondrial respiratory chain functions in BAT.

## Materials and Methods

### Chemicals

All chemicals were purchased from Sigma Aldrich unless otherwise noted.

### Animals

All mice were males between 8 and12 weeks of age at the time of sacrifice. They were fed standard chow and water *ad libitum*, and maintained on a light dark cycle of 12hr in a room maintained at 23 +/− 2°C. Wild type, UCP1KO, UCP3KO, and UCP1/3KO animals were on a C57B6/J background. UCP1KO mice were a gift from Dr. Leslie Kozak (Maine Medical Research Institute, Scarborough, ME, USA); UCP3KO were a gift from Mark Reitman, formerly of the National Institutes of Health. UCPDK (UCP1/UCP3 double knockout animals) mice were generated in house, by breeding heterozygous the UCP1KO mouse with the heterozygous UCP3KO mouse. Every five generations, animals from all genotypes were backcrossed with an in-house C57B6/J colony, to prevent genetic drift. Animals were euthanized via isoflurane inhalation followed by cervical dislocation. The University of Texas IACUC approved all animal experiments.

### Mitochondrial isolation

Mitochondria were isolated as described in [16]. Briefly, intrascapular BAT was harvested from 2-5 mice 8-12 week old male mice, rinsed in ice cold PBS, and cleaned of surrounding white adipose tissue. BAT was then minced with fine scissors, and homogenized in 250 mM sucrose. The homogenate was filtered through a 40uM cell strainer and centrifuged at 8,500 × *g*. The fat layer and supernant were removed; the pellet was resuspended in sucrose and centrifuged at 800 × *g*. Resultant supernant was centrifuged, and the pellet was resuspended in sucrose supplemented with 0.3% fatty acid free BSA, and pelleted at 8,500 × *g*. The resulting pellet was resuspended in 200ul of 250mM sucrose, and protein was measured with use of an BCA assay (Pierce). All steps were performed at 4°C, or on ice - unless otherwise noted. After preparation, mitochondria were either used directly for oxygen consumption measurements, or was frozen at −80°C for further analysis.

### Mitochondrial respiration

Isolated mitochondria (0.25-0.5mg/ml) in respiration buffer 0.6mL (0.1% BSA, 125mM sucrose, 20mM TES, 1mM EDTA, 4mM KP_i_, 2mM MgCl_2_ pH 7.2) were added to a continuously mixed chamber kept at 37°C (Hanstech); respiration was monitored for 15-30 seconds, after which pyruvate (5mM) and malate (2.5mM), GDP (1mM), ADP (1mM), Oligomycin (1.3ug/mL), and FCCP (0.5-2uM) were added sequentially. In a separate run, titration of respiratory activity was induced by oleic acid (50% EtOH) after mitochondrial respiration had been inhibited by 1mM GDP and 1.3ug/mL oligomycin. Oxygen consumption was measured every second, until the experiment was completed. All mitochondrial respiratory chain inhibitors were solubilized in 95% EtOH unless otherwise noted, and total EtOH up to 0.1% was found to have no effect on respiration.

### Mitochondrial complex activity assays

Mitochondrial complex assays were run essentially as described in [17]. Frozen mitochondrial aliquots were thawed on ice, then freeze-thawed an additional two times using a dry ice bath. Prepared mitochondria were used at a concentration of 4ug/ml (CS, CI, CIV) or 5ug/ml (CII, CI-III, CII-CIII). Absorbance was read on a Cary 40000 UV-Vis spectrometer at 37°C. Rates were calculated in Microsoft Excel, using linear portions of the activity. All complex activities were normalized to citrate synthase activity.

### Western blotting

Fresh mitochondrial lysates were prepared and diluted (2ug/ul) in sample buffer, and incubated for 5 minutes at 95°C. Mitochondrial proteins were loaded and run on a Tris-glycine gel system and transferred to nitrocellulose membranes, which were then incubated overnight with 5% milk in TBS-tween, followed by incubation with primary antibodies, and then for an additional hour with secondary antibodies, washed, and developed using ECL (Thermo-scientific).

### High resolution Clear Native PAGE

Pelleted mitochondria (500ug) were resuspended in 50ul of buffer (50mM NaCl, 50 mM imidazole, 2mM 6-aminohexanoic acid, 1mM EDTA, pH 7) and solubilized with digitonin (2g/g at 20%). Supernant was cleared by spinning at 20000 × *g* for 20 minutes. Glycerol was added (50% 2.5ul), and 10ul (~75ug) was loaded on a 3-12% Bis-Tris native gel. The Anode buffer was comprised of (25mM imidazole pH 7) and the cathode buffer (50 mm Tricine, 7.5 mm imidazole, 0.01% DDM pH 7). Gels were run until the MW marker reached the end of the gel, then subjected to sequential activity assays. <u>Complex I-II activity</u>: Gels were incubated for 10 minutes in Complex I buffer (2.5mg/ml nitrotetrazolium blue, 1 mg/ml NADH in 5mM Tris/HCl), transferred to 5mM Tris/HCl buffer, scanned, and then incubated for an additional 20 minutes in Complex II buffer (2.5mg/ml nitrotetrazolium blue, 20mM Succinate, 2uM phenazine methanosulfate, 5mM Tris/HCl) and imaged. <u>Complex III-IV activity</u>: Gels were incubated for 30 minutes in Complex III buffer (.5mg/ml diaminobenzidine, 50mM sodium phosphate, pH 7.2), scanned imaged, and placed back in Complex III buffer; cytochrome c was added to a 50uM final concentration, gels were incubated for another 30 minutes, and imaged again.

### Statistics

For all experiments a one-way ANOVA followed by a *Fisher LSD* test was used to compare multiple genotypes, with significance set *a priori* at p <.05.

## Results and Discussion

### UCP1 is a dominant regulator of brown adipose tissue respiratory chain function, expression, and assembly in BAT mitochondria

To better understand how UCP1/3 affects mitochondrial oxidation, we measured the activity of respiratory chains. We found that the loss of UCP1 in the presence or absence of UCP3 decreased all respiratory complex activities that were measured (Figure 1). The loss of UCP1 severely impacted complex I activity in both UCP1 and UCPDK mice, with only ~20% of WT activity remaining (In contrast, complex II activity is ~50% of WT activity). Complex IV activity was also decreased in the absence of UCP1, to about 60% of WT activity. Notably, complex IV activity was significantly increased in the UCP3KO mitochondria (Figure 1C), and in agreement with the report by Liebig *et al.* [18].

**Figure 1.**
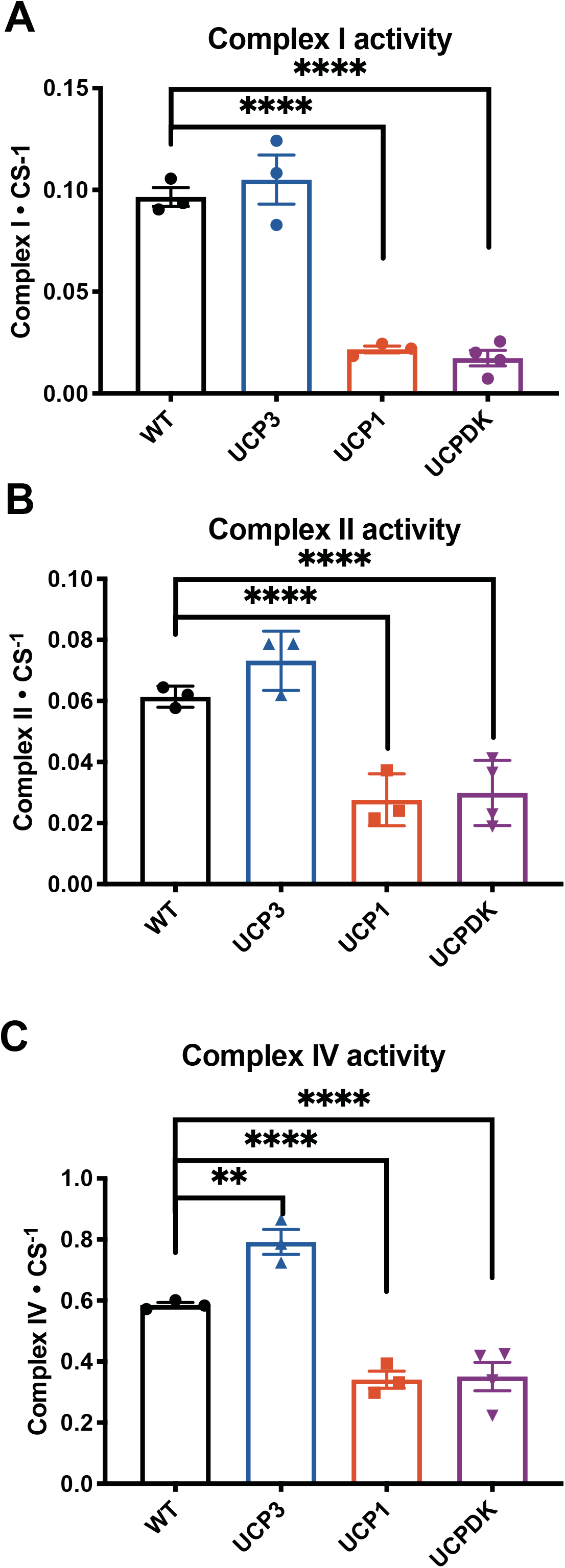
Loss of UCP1 is the dominant regulator of mitochondrial respiratory chain enzyme activity. (A) Complex I activity (B) Complex II activity and (C) Complex IV activity were each controlled for total mitochondria, using citrate synthase activity. Freeze-thawed mitochondria from WT (n=3) UCP1KO (n=3) UCP3KO (n=3) and UCPDK(n=4). *\*= p< .05, **=p<.001, ***= p<.0001, ****= p<.00001*.

To explore the decrease of respiratory chain activity, we determined protein abundance via western blotting, and found that the loss of UCP1 decreased Ndufa9 (complex I), Uqcrc2 (complex III), and Coxiv (complex IV) protein abundance in mitochondria (Figure 2A). Coxiv expression was also decreased in the double knockout background. However, expression of Sdha (complex II) was not affected, despite a reduction in its activity (Figure 1B, 2A). Neither the expression of Oscp (complex V) nor of Mdh2(TCA cycle enzyme) was changed in any of the genotypes (Figure 2). Because these enzymes only represent a selected few of the many components of each respiratory complex, we investigated how both UCP’s influence the formation of respiratory chain complex assembly in BAT. Using activity-based high resolution clear native PAGE, we confirmed that in the absence of UCP1, complex I, complex III, and complex IV are depleted and disorganized (Figure 2C), and complex III occupancy was decreased in supercomplexes (Figure 3C). However, no additional loss of mitochondrial complexes was detected in the UCPDK mice when compared with mice in which only UCP1 was depleted (Figure 3C).

**Figure 2.**
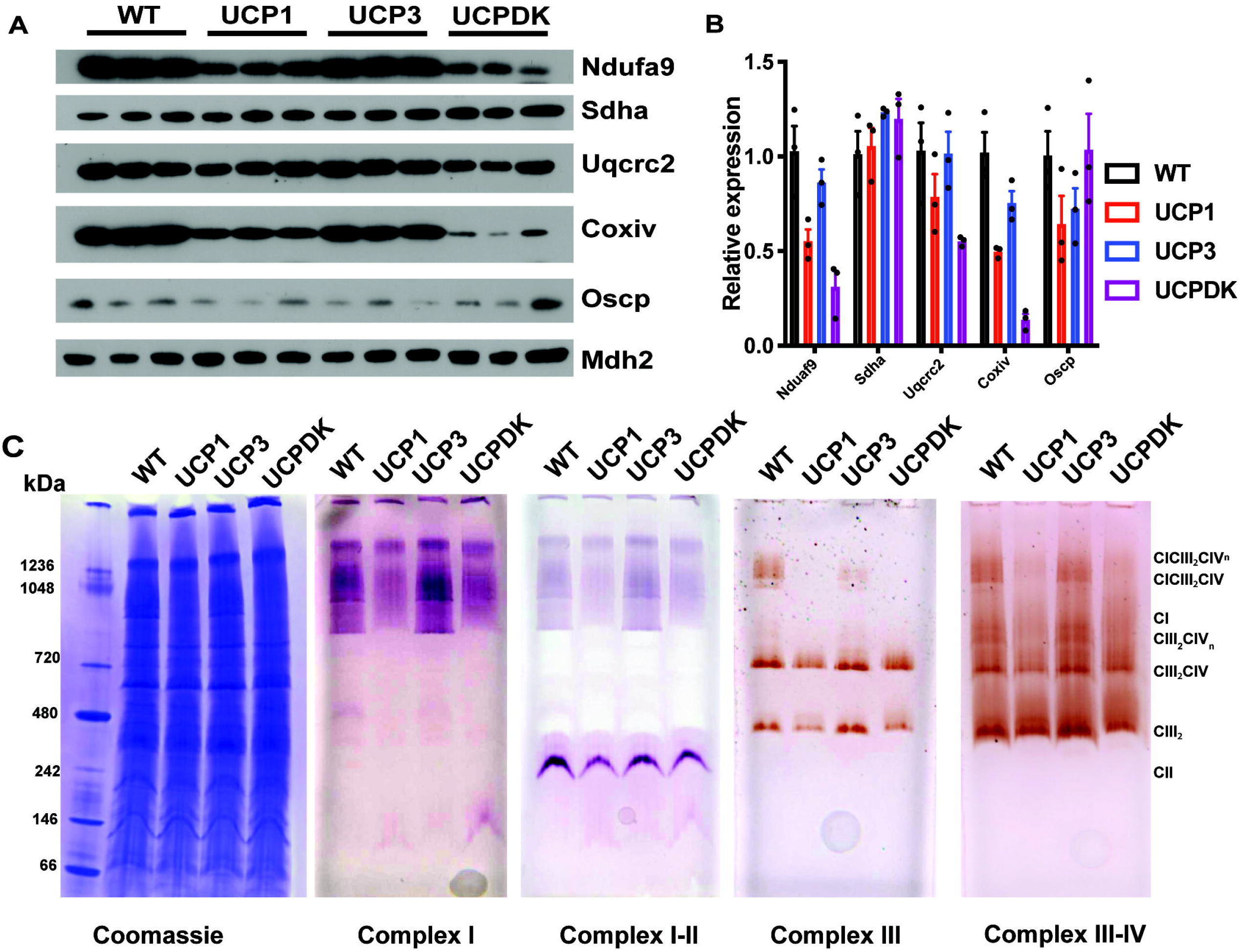
Loss of UCP1 is the dominant regulator of respiratory chain protein expression and organization in brown adipose tissue mitochondria. (A) Western blots of mitochondrial extracts from indicated genotypes. Five micrograms of mitochondrial protein from three separate mitochondrial preparations per genotype. (B) Densitometry from data in *A* was quantified using ImageJ, and was controlled for Mdh2 expression. N=3 per genotype. (C) Representative high resolution clear native PAGE analysis of mitochondria from indicated genotypes.

**Figure 3.**
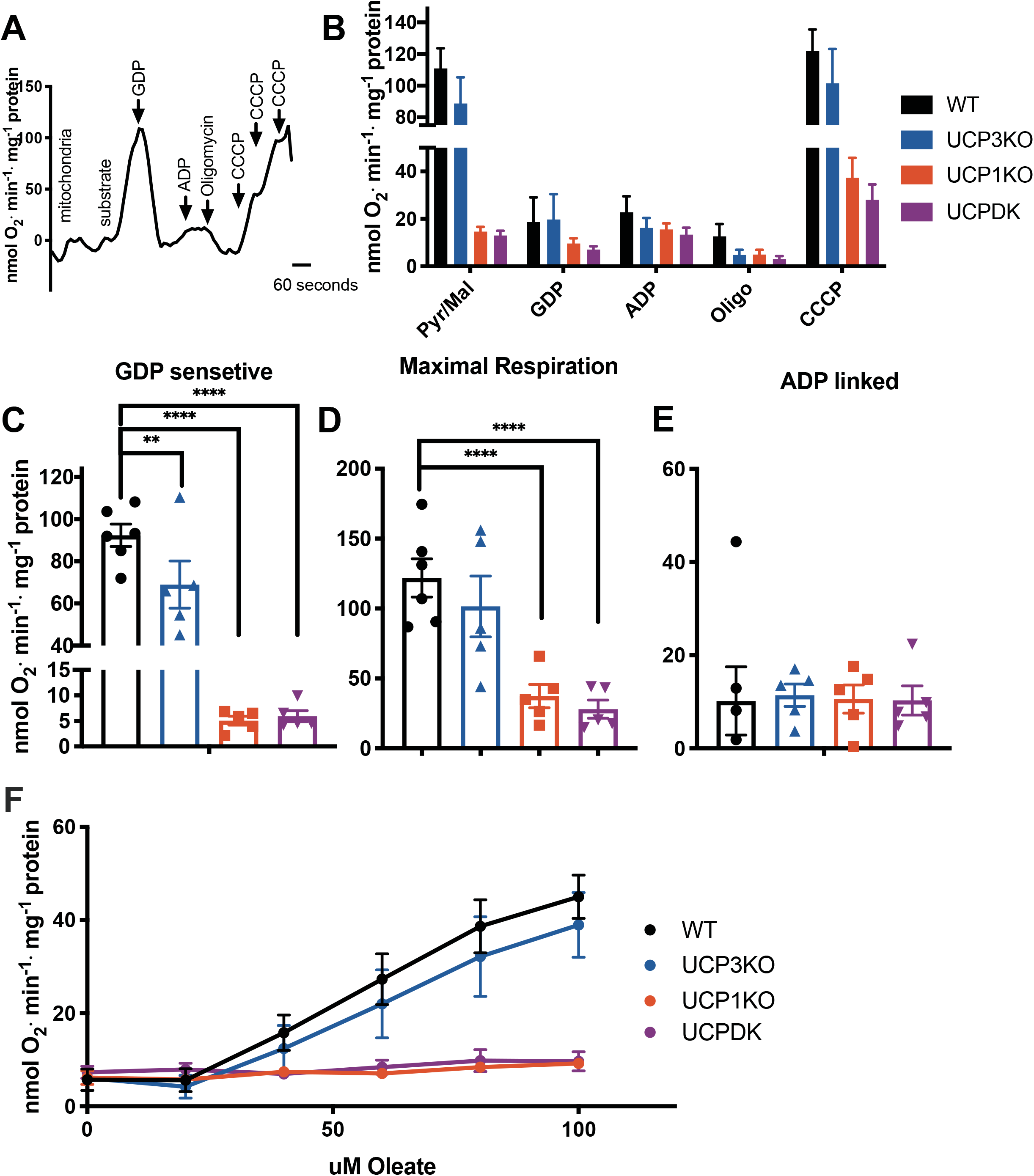
Regulation of brown adipose tissue mitochondrial respiration by uncoupling proteins 1 and 3. (A) Scheme for mitochondrial respiration assays, (B) Mitochondrial respiration from 5-6 different preparations per genotype, (C) GDP-sensitive respiration, (D) Maximal respiration, (E) ADP-linked respiration, and (F) Oleate induced respiration. (A-E) n= 5-6 per genotype *\*= p< .05, **=p<.001, ***= p<.0001, ****= p<.00001*.

Together, these data unambiguously demonstrate that Ucp1 is necessary for thermogenically competent brown adipose mitochondria. Our data strongly support those of Kazak *et al.* We also expand their finding that Ucp1 is essential for the functioning of the brown fat respiratory chain at 4C, to include mice that are raised under standard housing conditions[19]. The loss of Ucp1 extensively alters the BAT proteome, and highlight that UCP3 cannot compensate for the loss of Upc1, even though Ucp3 is expressed in this model [9].

The discrepancies in Sdha protein abundance and complex II activity between WT and UCP1-null animals points to the existence of a regulatory function that does not allow electron transfer between the matrix-exposed Sdha-Sdhb complex and the membrane-embedded Sdhc-Sdhd complex. Confirmation of increased complex IV activity, despite similar protein abundance and complex integrity in the UCP3-null mouse and the Ucp3-null hamster, as shown by Liebig’s work, suggests that this phenomenon reflects a mechanism wherein an increase complex IV activity may compensate for the loss of Ucp3 function.

### Loss of UCP3 decreases GDP sensitive respiration, but does not affect FFA-induced respiration in BAT mitochondria

As mitochondria are an integrated system, we used high-resolution respirometry to further probe the bioenergetic function of UCP3 in brown adipose mitochondria. The protocol outlined in Figure 4A utilizes continuous addition of substrates and inhibitors, to elucidate GDP-sensitive (respiration after addition of substrate minus respiration after GDP addition), ADP-linked (respiration after addition of ADP minus respiration after oligomycin), maximal respiration (CCCP induced) (Fig. 3A) [20]. In agreement with the complex activity data, loss of UCP1 was found to be the dominant regulator of respiration (Fig. 3B-D). Notably, the loss of UCP3 only (i.e., with UCP1 present) led to a decrease in GDP-sensitive respiration (Fig. 3C). ADP-linked respiration was equal across all genotypes, despite the significant decrease in total respiratory capacity (Fig. 3E). We found no significant difference in free fatty acid-mediated uncoupling in the presence or absence of UCP3(Fig. 3F).

The finding that ADP-linked respiratory capacity is the same between the UCP1-null and WT mitochondria is critical: it suggests that the recently discovered ATP-linked thermogenic pathways remain unaffected by the loss of UCP1. This finding means that proposed futile cycles that require ATP production (phospho-creatine and calcium) can run in the absence of UCP1 in BAT[21,22]. However, total respiratory capacity in this tissue is likely compromised, and heat production would be severely impaired relative t WT animals.

Previous reports suggest that the role of Ucp3 in BAT is negligible. However, those studies and the data presented in this report have critical differences. Gong *et al.* used a mouse model on a mixed background that is not highly reliant on BAT thermogenesis[23]. Also, they employed succinate as the oxidative substrate - given that succinate is not compatible with maximal oxidation in BAT mitochondria, their results may have underestimated the function of UCP3[16]. Additionally, when both UCP3 expression and adrenergic tone in BAT were low, [16] they housed their mice at thermoneutrality [16]. And Liebig *et al.* used a mutant hamster with BAT-specific ablation of UCP3, and also found no significant effect on mitochondrial oxidation when respiration was driven with glycerol-3-phosphate (G3P). In contrast, here we have used pyruvate and malate, a complex I-linked substrate that requires the transport of both substrates across the inner membrane.

Discrepancies between our findings and earlier reports may be explained by UCP3’s hypothesized transport function, through which it increases the availability of substrates pyruvate or malate during maximal UCP1 activation[11]. UCP2, which shares a 72% homology with UCP3, directly transports C4 metabolites, and controls glycolysis as well as glutamine oxidation in hepatocytes [24]. A recent study has demonstrated biochemical differences between UCP1 and UCP3 that point to differing transport roles[25]. In light of existing literature, our data suggest that UCP3 complements UCP1’s uncoupling function, not as an uncoupler but as a solute transporter.

In summary, our works shows that UCP3 does have a role in mitochondrial oxidation in brown adipose tissue mitochondria. Future work will investigate the possible transport functions of this protein and its role in substrate oxidation more globally.

